# The methylome of the human frontal cortex across development

**DOI:** 10.1101/005504

**Authors:** Andrew E. Jaffe, Yuan Gao, Ran Tao, Thomas M. Hyde, Daniel R. Weinberger, Joel E. Kleinman

**Author notes:** + equally contributing authors (Daniel R. Weinberger, Joel E. Kleinman).

## Abstract

DNA methylation (DNAm) plays an important role in epigenetic regulation of gene expression, orchestrating tissue differentiation and development during all stages of mammalian life. This epigenetic control is especially important in the human brain, with extremely dynamic gene expression during fetal and infant life, and becomes progressively more stable at later periods of development. We characterized the epigenetic state of the developing and aging human frontal cortex in post-mortem tissue from 351 individuals across the lifespan using the Illumina 450k DNA methylation microarray. The largest changes in the methylome occur at birth at varying spatial resolutions – we identify 359,087 differentially methylated loci, which form 23,732 significant differentially methylated regions (DMRs). There were also 298 regions of long-range changes in DNAm, termed “blocks”, associated with birth that strongly overlap previously published colon cancer “blocks”. We then identify 55,439 DMRs associated with development and aging, of which 61.9% significantly associate with nearby gene expression levels. Lastly, we find enrichment of genomic loci of risk for schizophrenia and several other common diseases among these developmental DMRs. These data, integrated with existing genetic and transcriptomic data, create a rich genomic resource across brain development.

## Introduction

DNA methylation (DNAm) plays an important role in epigenetic regulation of gene expression, orchestrating tissue differentiation and development during fetal life, childhood, and adolescence, and guiding functional activity in adulthood. This epigenetic control is especially important in the human brain, where gene expression is extremely dynamic during fetal and infant life, and becomes progressively more stable at later periods of development (Colantuoni et al. 2011; Numata et al. 2012). Dysregulation of these precise and coordinated gene expression changes through epigenetic mechanisms may play a vital role in the pathogenesis of neurodevelopmental disorders, such as schizophrenia (SZ) (Waterland and Michels 2007; Jakovcevski and Akbarian 2012; Grayson and Guidotti 2013). Pathologically, these epigenetic changes, acting through gene expression, could disturb the formation of essential brain circuits, fitting into one prevailing set of hypotheses for the causes of schizophrenia, namely the “neurodevelopmental” hypotheses (Weinberger and Levitt 2011).

DNA methylation is an attractive epigenetic mechanism to study in post-mortem human brain tissue for better understanding neurodevelopmental disorders, for reasons beyond its association with mRNA expression levels (Irizarry et al. 2009). Exogenous factors have been associated with altering DNAm levels, both at specific loci and globally (averaged across all repeat elements), including changes in diet (Heijmans et al. 2008), and exposure to cigarette smoking (Breitling et al. 2011) and arsenic (Reichard et al. 2007). For example, although schizophrenia usually presents in the third decade of life, the pathological changes that lead to this disorder may precede the onset of illness by several decades. Extensive research implicates environmental variables in the development of schizophrenia, especially acting during fetal and perinatal life, including maternal stress and infections, obstetric complications, and maternal nutrition during pregnancy (Weinberger and Levitt 2011), including during the Dutch famine of 1944-1945 led to a spike in the number of cases of schizophrenia two decades later (Susser and Lin 1992). Many of these factors have previously been associated with altering DNA methylation levels (Cortessis et al. 2012; Relton and Davey Smith 2012). Lastly, several recent papers have explored the role of sequence variation on site- and region-specific DNA methylation (Schilling et al. 2009; Lienert et al. 2011). The DNA sequence itself plays a large role in the maintenance of DNAm (Bird 2011), providing one potential mechanism, namely changes in DNAm, for the clinical associations of single nucleotide polymorphism (SNPs) from large genome-wide association studies (GWAS) like the Psychiatric Genetics Consortium (Sullivan et al. 2012).

We generated epigenome-wide DNA methylation (DNAm) from postmortem dorsolateral prefrontal cortex (DLPFC) brain tissue from 351 non-psychiatric and medically “normal” subjects, including 35 second trimester fetal samples, using the Illumina HumanMethylation450 (“450k”) microarray to better characterize changes in DNAm across development and aging. This work extends previous DNAm maps of human frontal cortex by twenty-fold increased genomic coverage (485,000 versus 27,000 probes) in a much larger sample size (351 versus 108 samples) (Numata et al. 2012) creating a more comprehensive landscape of epigenetic development in the human brain. This platform improves functional genomics analyses by more likely associating epigenetic changes with gene expression changes via many probes in CpG island shores (Irizarry et al. 2009) and the ability to identify changes in DNAm on the region-, rather than probe-, level (Aryee et al. 2014). which better associate with gene expression. While previous efforts to comprehensively measure DNA methylation across the epigenome using whole genome bisulfite sequencing have identified many important features of brain development (Lister et al. 2013), we complement this work using a much larger sample at more continuous ages samples, albeit at lower genome-wide coverage, to obtain population-level spatial dynamics of DNA methylation across brain development. We have implemented novel statistical methods to identify regional and long range changes in DNAm to overcome some of the shortcoming of microarray data, and find extensive evidence of DNAm changes at birth. We then more formally define regions of developmental importance, termed “development DMRs”, that capture continuous patterns of age which strongly associate with the gene expression levels of nearby genes. Lastly, these developmental DMRs are enriched for clinical risk loci for schizophrenia and other common diseases identified by large multi-study meta-analyses. Our results suggest large-scale changes in the human brain methylome across development and aging with potentially functional consequences.

## Results

### Changes in the methylome at birth

First, we analyzed fetal (n=35) compared to non-fetal (n=316) samples (including newborns and children) to identify changes in DNAm associated with birth, at varying spatial scales. At the single probe level, the majority (N=359,087, or 78.5% of probes on the array) of assayed CpGs were significantly differentially methylated (at p < 0.05), suggesting a vastly different epigenetic landscape of the prefrontal cortex during fetal compared with postnatal life (Figure 1A). These differentially methylated CpGs were classified into differentially methylated regions (DMRs) based on a “bump hunting” approach (Jaffe et al. 2012), resulting in 23,732 statistically significant DMRs (Figure 1B). Lastly, we identified 298 regions of long-range differential methylation (Figure 1C), termed “blocks” (Hansen et al. 2011), using an approach adapted to the Illumina 450k (Aryee et al. 2014) from whole genome bisulfite sequencing (WGBS) data first utilized in comparing colon cancer to normal tissue (Supplementary Table 1). We found significant overlap of fetal-associated prefrontal cortex blocks with these cancer blocks (273 of the 298; 91.6%), including consistent directionality – fetal samples were almost exclusively hypomethylated compared to adult samples, which mirrored the hypomethylated cancer blocks (85.0% of overlapping blocks, Supplementary Figure 1), suggesting these blocks may represent more general developmental and/or proliferative phenomena.

**Figure 1:**
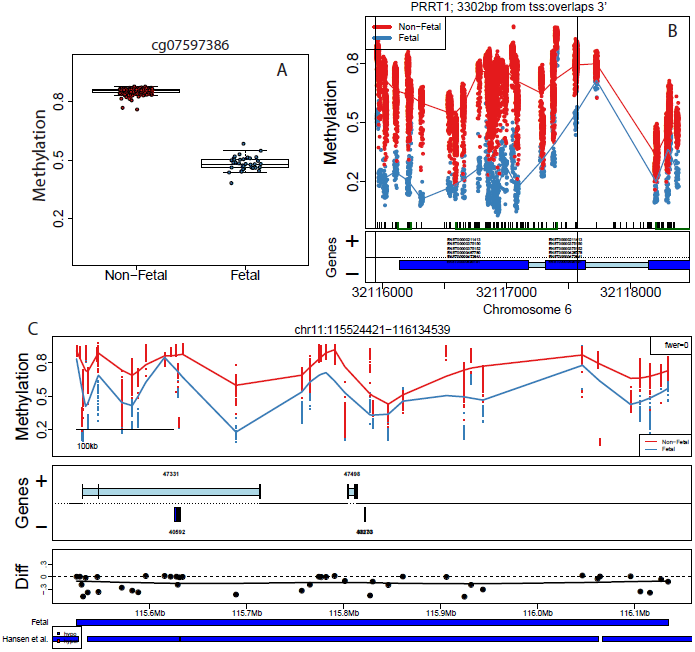
fetal versus non-fetal differentially methylated loci at three spatial resolutions. Examples of significant (A) differentially methylated probes representing local changes, (B) differentially methylated regions (DMRs) representing regional differences and (C) methylation blocks representing long range changes, are shown comparing fetal versus non-fetal samples. Proportion methylation is shown on the y-axis of each respective top-most panel. Gene annotation panels in (B) and (C) are based on Ensembl annotation – dark blue represents exons and light blue represents introns. Ensembl transcripts annotate genes in (B) and Entrez Gene IDs annotate genes in (C). The third panel in (C) depicts the average difference in DNAm between groups at each probe group and the final panel shows the overlap between cancer hypomethylation blocks from Hansen et al. (2011). Blue = fetal samples; red = non-fetal samples.

These strong global effects may reflect composition changes of the underlying brain tissue comparing fetal frontal cortex (predominantly neurons and neuronal precursors) and adult prefrontal cortex (mixture of neurons and glia). To address this potential issue, we utilized the publically available sorted data from Guintivano et al. (2013) (see Methods) to assess statistical enrichment among the resulting fetal DMR list compared to adult neuronal versus non-neuronal differences. We identified strong significant enrichment among those DMRs called significant between datasets (odds ratios for enrichment greater than 40, corresponding to p< 10^−100^) with approximately 87% of cell-type DMRs containing fetal DMRs. These results suggest a shifting epigenetic landscape driven by cellular composition shifts with site-specific changes within pure cellular populations.

We also explored our results in the context of the recently-published differentially CpG-methylation regions reported by Lister et al. (2013) in WGBS data from one fetal compared to two flow-sorted (again into NeuN+ and NeuN-) adult frontal cortices. Of the 267,799 regions across four comparisons (NeuN+/- and hyper/hypomethylated), there were 51,211 regions with coverage on the Illumina 450k, of which 39,697 (77.5%) were significantly methylated in the fetal versus postnatal comparison at p < 0.05 and directionally consistent (hyper- or hypo-methylated in the same direction across datasets). The hypermethylated DMRs within the NeuN- sample had the greatest differences in our data, which, given the design of the 450k (mostly unmethylated loci) and composition change (gain in NeuN- fraction) across development, suggests an overall comparability across datasets. However, there were 5,649 significant regions (11.0%), with opposite directionality of methylation and another 5,889 regions (11.4%) not significantly different by life stage. We also identified thousands of DMRs in our larger sample that were not identified as differentially methylated in the WGBS data, likely due to the increase in power of our much larger number of biological samples. Biological variability across samples even within specific age ranges can therefore be large in certain regions of the epigenome, and these DNA methylation maps of bulk tissue and within sorted cell types requires large samples for adequate power.

### Developmental DMRs associate with nearby gene expression levels

We further explored more subtle patterns of DNAm across the lifespan and identified 55,439 significant differentially methylated regions (DMRs) associated with development and aging (Supplementary Methods) beyond fetal versus non-fetal life stages, representing 126,866 probes on the microarray (27.8%) (Figure 2, Supplementary Table 2). While the majority of the DMRs were driven by large fetal versus non-fetal differences, infant and child samples were directionally consistent, oftentimes with DNAm levels intermediate between fetal and adult samples (Supplementary Figure 2). An important observation was that among the post-natal samples, the estimated neuronal composition (via empirical NeuN+ proportion) went down across the lifespan, on average (p=3.22×10^−4^), reflecting the shifting composition of the young and developing brain, as glial cells are born and migrate (Supplementary Figure 3). We similarly see a strong enrichment of NeuN DMRs among these more developmental DMRs (p < 10^−100^), suggesting again that brain development epigenetically reflects shifting cellular populations combined with changes in site-specific DNAm within cell populations.

**Figure 2:**
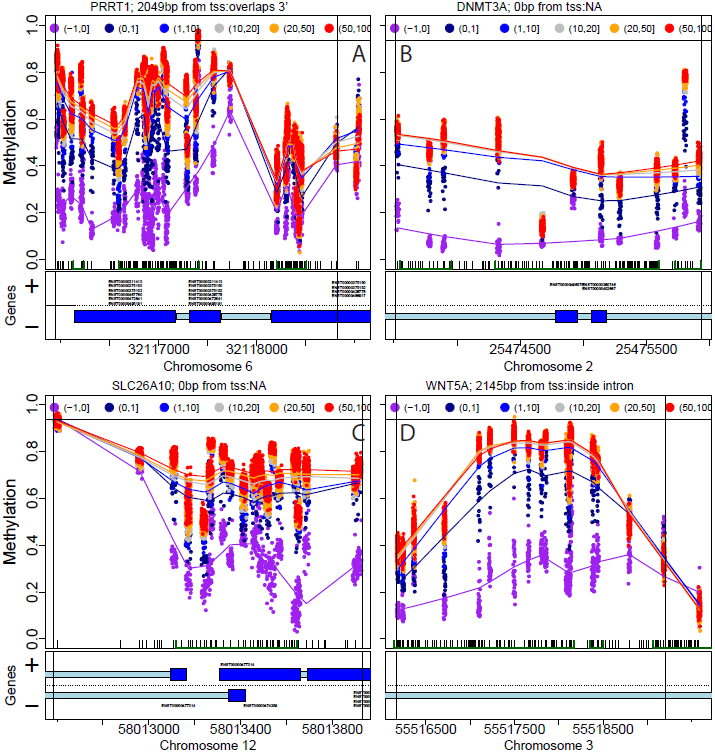
methylation plots for four example differentially methylated regions (DMRs) for development. Panel A: *PRRT1,* Panel B: *DNMT3A,* Panel C: *SLC26A10,* Panel D: *WNT5A.* Top two-thirds of panels depict individual methylation levels at each probe by genomic position, with colored lines reflecting the average methylation curve for samples binned by age group as shown in the legend. Tick marks show the location of CpG dinucleotides and green bars indicate annotated CpG islands. The bottom panel shows the location of Ensembl annotated genes (dark blue: exons annotated by Ensembl transcripts, light blue: introns, + and - represent the direction of the gene). Vertical lines represent boundaries of the DMR, which can span the entire probe group.

We functionally characterized these “developmental DMRs" first by mapping each DMR to its nearest gene. Among the 55,439 significant DMRs, more than half lie within gene bodies (53.6%), 7.0% lie in gene promoters, and another 10.5% and 10.7% were upstream and downstream of genes respectively. A subset of the samples had corresponding gene expression data (N=260), publicly available from the Gene Expression Omnibus (GEO, available at GSE30272; see Methods section) (Colantuoni et al. 2011). The correlation between DNAm (average level of DNAm across the region) of each DMR and its corresponding nearest gene’s expression was assessed across the lifespan: 61.9% of DMRs (N=31,822) were significantly correlated with neighboring gene expression (at p < 10^−10^) (Figure 3). Statistically significant DMRs were highly associated with gene expression (p < 10^−100^), and more specifically, the strength of region-level differential methylation strongly predicts the DNAm-gene expression association (spearman correlation = 0.31, p < 10^−100^), reinforcing the notion that regional DNAm levels in brain correlate strongly with gene expression levels as has been also observed in other tissue and cancer methylation differences (Doi et al. 2009). Interestingly, among the 61.9% of significant DMRs associated with gene expression, many of these DNAm-expression pairs are positively correlated (46.4% of significant DNAm-expression pairs).

**Figure 3:**
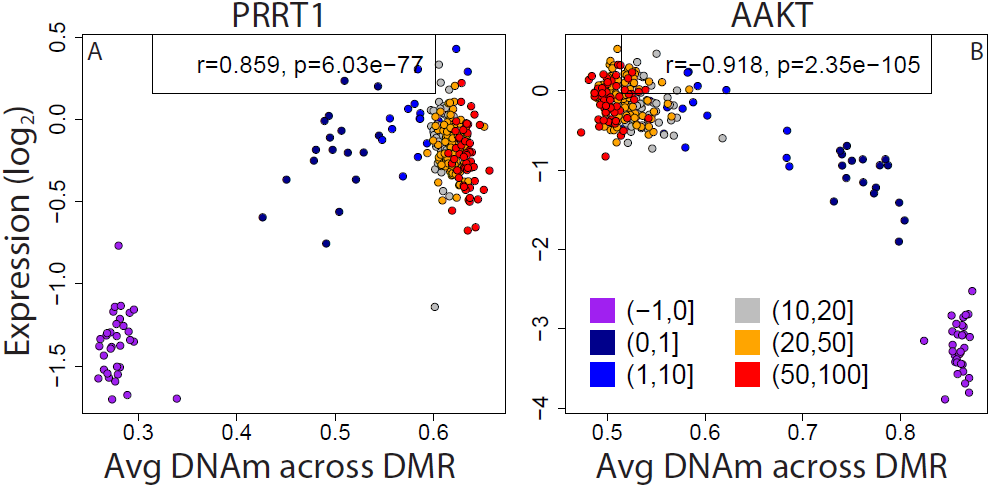
correlation between DNA methylation (DNAm) and gene expression for (A) *PRRT1* and (B) *AAKT.* Gene expression on the log_2_ scale, representing the fold change between each sample and a pool of many samples from across the lifespan, is plotted against the average DNAm for each sample within the significant DMR. Color indicates sample age, and the Pearson correlation with corresponding p-value is shown in the top legend.

### Developmental DMRs are enriched for expression quantitative trait loci (eOTLs)

We then asked whether DNA methylation mediates the effect of genetic variation on gene expression – each sample with gene expression data also has corresponding genome-wide single nucleotide polymorphism (SNP) data, imputed up to 6 million SNPs per sample. We identified 204,380 significant cis eQTLs in these data (at FDR < 1%, corresponding to p < 6.9×10^−5^), with linkage-disequilibrium blocks corresponding to 5,335 unique probes and 4,193 unique genes. We then identified the single CpG with the strongest association to each gene’s expression (within gene body +/- lokb). We observed stronger association between gene expression and DNA methylation among those genes (n=5,335) that were associated with a nearby SNP (i.e. an eQTL; p = 9.18×10^−39^), suggesting that DNA methylation levels may indeed mediate the effect of genetic variation on gene expression.

### Developmental DMRs are enriched for schizophrenia genetic risk loci

Lastly, we identified significant association between genomic loci implicated in schizophrenia risk from the latest Psychiatric Genetics Consortium (PGC) data release (Ripke et al. 2013) and our developmentally significant DMRs. There was significant overlap (Table 1, p=0.0019 – 0.047) in the locations of developmental DMRs (at FWER < 5%) and clinical risk regions (starting at loci with association p < 10^−4^ through genome-wide significance 5×10^−8^), suggesting these regions of schizophrenia risk are important for brain development potentially through differential DNA methylation between fetal life and infanthood. These results are consistent with prevailing hypotheses about the role of early brain development and risk for schizophrenia (Weinberger and Levitt 2011). In additional analyses, we note that significant enrichments are not unique to schizophrenia – we find significant enrichment for genome-wide significant loci (at p < 5×10^−8^, via Supplementary Table 2) associated with inflammatory bowel disease (p=1.9×10^−8^) and ulcerative colitis (p=1.12×10^−4^) but not Crohn’s disease (p=0.25) (Jostins et al. 2012), and enrichment of type 2 diabetes loci that failed to reach genome-wide significance (enrichment p < 0.05 for GWAS p-values less than 5×10^−7^) (Morris et al. 2012). These developmental DMRs may therefore reflect regions in the epigenome that are dynamic across other tissues – 52% of the developmental DMRs overlapped “dynamic DMRs” across 30 human cell and tissue types identified by Ziller et al. (2013) (compared to 41% of probes groups on the Illumina 450k).

**Table l:** Enrichment of schizophrenia and diabetes genetic risk regions

|  | **Schizophrenia** |  | **Type 2 Diabetes** |  |
| --- | --- | --- | --- | --- |
| **GWAS Cutoff** | **# gwas sig** | **devDMR enrichment p-value** | **# gwas sig** | **devDMR enrichment p-value** |
| p<1e-04 | 867 | 0.047 | 381 | 0.003 |
| p<5e-05 | 541 | 0.016 | 302 | 0.029 |
| p<1e-05 | 199 | 0.0019 | 210 | 0.00066 |
| p<5e-06 | 131 | 0.034 | 65 | 0.0016 |
| p<1e-06 | 62 | 0.0028 | 37 | 0.0029 |
| p<5e-07 | 41 | 0.002 | 34 | 0.0075 |
| p<1e-07 | 16 | 0.0095 | 17 | 0.52 |
| p<5e-08 | 12 | 0.011 | 16 | 0.45 |
| p<1e-08 | 6 | 0.068 | 13 | 0.39 |
| p<5e-09 | 6 | 0.068 | 13 | 0.39 |
| p<1e-09 | 5 | 0.068 | 10 | 1 |
| p<5e-10 | 4 | 0.068 | 10 | 1 |
| p<1e-10 | 3 | 0.063 | 9 | 1 |

## Discussion

In summary, we explore DNA methylation changes across the lifespan in the human DLPFC in the largest epigenetic study of brain tissue to date, and identify widespread change in the epigenome occurring at birth at local, regional, and long-range spatial resolutions. These large changes in DNAm likely represent in part shifts in neuronal composition across the lifespan, with site-specific changes within pure neuronal and non-neuronal cellular populations, and correspond to strong changes in gene expression profiles in the majority of significant differentially methylated regions (DMRs) associated with aging. Furthermore we find significant enrichment of these developmental DMRs with genes associated with both genetic control of gene expression and clinical risk for schizophrenia and other common complex genetic disorders. These raw data are publicly available on GEO (Edgar et al. 2002) under accession GSExxxx, and user-friendly data will be made available from our BrainCloud desktop application (http://braincloud.jhmi.edu), both containing information to link these DNAm samples to existing publicly available data on gene expression (GSE30272) and genetic variation (phs000417.v1.p1) to create a rich resource of genomic data across brain development.

Deviations from these essential DNAm developmental trajectories during critical windows of development from conception to young adulthood may interfere with the carefully coordinated temporal and spatial dynamics of gene expression through a combination of genetic and epigenetic factors (Abdolmaleky et al. 2004; Mill et al. 2008; Jakovcevski and Akbarian 2012; Grayson and Guidotti 2013). These characterized patterns of DNA methylation at specific genomic loci in the developing brain therefore has clear implications for better understanding the role that epigenetics plays in neurodevelopmental disorders. Mechanistically, these changes in DNAm may be the crucial link by which environmental events amplify the effects of genetic variations in increasing liability towards illness. The development of novel treatment strategies for these disorders is dependent upon understanding the molecular pathways that increase liability towards illness. Furthermore, these patterns of DNA methylation at the population level during the first three decades of life may elicit a better understanding how genetic variation interacts with environmental factors in altering risk for illness.

## Methods and Materials

### Studu samples

Brain specimens were donated through the Offices of the Chief Medical Examiners of the District of Columbia and of the Commonwealth of Virginia, Northern District to the NIMH Brain Tissue Collection at the National Institutes of Health in Bethesda, MD, according to NIH Institutional Review Board guidelines (Protocol #90-M-0142). Audiotaped informed consent was obtained from legal next-of-kin on every case. Details of the donation process are described elsewhere (Deep-Soboslay et al. 2005; Lipska et al. 2006). Additional specimens, including the 35 second-trimester fetal brain tissue samples, were obtained via a Material Transfer Agreement with the National Institute of Child Health and Human Development Brain and Tissue Bank. All postnatal non-psychiatric control cases (N=316) were free from psychiatric and/or neurologic diagnoses and substance abuse according to DSM-IV. Every control case had toxicology screening to exclude for acute drug and alcohol intoxication/use at time of death, and all fetal tissue was also screened for possible in utero drug exposure.

### Tissue Processing

All specimens were flash-frozen, and screened for macro- and microscopic neuropathological abnormalities, as previously described (Lipska et al. 2006). All specimens with significant evidence of neurological disorders, infarcts or other cerebrovascular abnormalities were excluded from study. Brain pH was measured, and postmortem interval (PMI, in hours) was calculated for every sample. Postmortem tissue homogenates of the prefrontal cortex (dorsolateral prefrontal cortex, DLPFC, BA46/9) were obtained from all subjects. Genomic DNA was extracted from 100 mg of pulverized dorsolateral prefrontal cortex (DLPFC) tissue with the phenol-chloroform method. Bisulfite conversion of 600 ng genomic DNA was performed with the EZ DNA methylation kit (Zymo Research).

### DNA Methylation Microarray

DNA methylation was assessed using the Illumina HumanMethylation450 (“450k”) microarray, which measures CpG methylation across >485,000 probes covering 99% of RefSeq gene promoters (Sandoval et al. 2011). Arrays were run following the manufacturer’s protocols. A percentage of the samples were run in duplicate across multiple processing plates to assess technical variability related to DNA extraction and bisulfite conversion. A total of 466 microarrays were scanned on 351 unique subjects.

### Data Processing and Normalization

Red and green channel intensity files were obtained for each sample in the “idat” file format. These files were processed and normalized using the *minfi* Bioconductor package in R (Aryee et al. 2014). Red and green intensities were mapped to the M(eth) and U(nmeth) channels, and the average intensity for these channels were used to check for low quality samples (o samples were dropped). Intensities from the sex chromosomes were used to predict sex, and we dropped 5 samples that had predicted sex different from its recorded value (indicating potential sample swaps). Then, the M and U channels were subsequently aeross-sample quantile normalized using an approach developed by Touleimat and Tost (2012). Briefly, this approach forces the distribution of type I and type II to be the same by first quantile normalizing the type II probes across samples and then interpolating a reference distribution to which the type I probes are normalized, stratified by region (e.g. promoter, shore, island, shelf). We found that across-sample normalization like quantile best minimized the variability between replicates. We retained a single array in the case of duplicates by choosing the sample that had the closest quality profile to all other arrays.

Flow-sorted samples from Guintivano et al. (2013) were included in this quantile normalization procedure via the FlowSorted.DLPFC.450k Bioconductor package to make the data more comparable in cellular composition estimation and differential methylation analysis (Jaffe and Irizarry 2014). In both datasets, probes on the sex chromosomes were dropped (which are difficult to normalize), as were probes annotated with single nucleotide polymorphisms (SNPs) at the target CpG or single base extension (SBE) site according to dbSNP137 with minor allele frequency > 1%, leaving 456,655 autosomal probes for analysis.

### Composition Estimation

We estimated the neuronal composition of each sample by simplifying the regression calibration approach developed for blood data (Houseman et al. 2012) using the Guintivano et al. (2013) flow sorted data at training data for the model. We selected 100 hyper- and hypo-methylated probes, and used linear regression to predict the relative proportion of neuronal cells in our heterogeneous DLPFC tissue. This statistical approach for neuronal composition estimation was accurate in fetal brain – there were significant differences in composition between fetal (27.3% NeuN+) and non-fetal (32.8% NeuN+) samples (p=2.33×10^−11^), but the directionality of the effect was antithesis to the underlying biology - we expect these brains to be composed almost entirely of neurons and/or neuron precursors, resulting in fetal estimates greater than non-fetal estimates (Supplementary Figure 3).

### Statistical Analyses for Differential Methylation

For the fetal versus non-fetal DLPFC analyses, single CpG analysis was performed using a t-test at every probe. Regional analysis to find differentially methylated regions (DMRs) and “block finding” were performed using the *minfi* R package (Hansen and Aryee 2013) using the *bumphunterEngine* and *blockFinder* functions, respectively, each with 1000 permutations and a cutoff of 0.1 (corresponding to a minimum 10% change in DNAm associated with birth). These analyses were all univariate given the very strong fetal effect. The NeuN+ versus NeuN− DMRs in the Guintivano et al. (2013) dataset were found using the same “bumphunting” procedure.

The developmental DMRs that allowed age to be modeled continuously were found using a modified version of the “bumphunting” algorithm (Jaffe et al. 2012). We fit a linear spline at every base, with knots at birth, then 1,10, 20, and 50 years of age – there was also an offset at birth because there were no samples in the third trimester. We then ran surrogate variable analysis (SVA) (Leek and Storey 2007) to model potential plate/”batch” effects. At each probe we computed an F-statistic comparing a statistical model containing the age spline plus estimated surrogate variables compared to a model containing just the surrogate variables. We then smoothed these f-statistics and thresholded within predefined probe groups using a single-CpG F-statistic p-value < 10^−20^ to identify DMRs. Statistical significance was assessed using the linear regression bootstrap which permutes the residuals around a regression fit 1000 times (Jaffe et al. 2012), and controlled for a family-wise error rate of 5%.

### Statistical Analyses for Gene Expression Correlation

Raw gene expression two-color microarray intensity data (available at GSE30272) were loess-normalized as previously described (Colantuoni et al. 2011) and matched to samples that had DNAm data (n=260). Surrogate variable analysis (SVA) was used with the age spline model described above to protect age effects and reduce observed technical processing plate and tissue quality effects in the data. At each developmental DMR, the average DNAm across that DMR in each sample was compared to the expression of the nearest gene, with matching based on gene symbol in the gene expression microarray annotation. If multiple expression probes existed for the gene, the probe most absolutely correlated with DNAm was retained. Pearson correlation was used to assess the relationship between DNAm and gene expression.

### Genotype data and eQTL analysis

DNA for genotyping was obtained from the cerebella of samples in the collection and performed with either the Illumina Human Hap 650V3 or lM Duo V3 BeadArrays as previously described (Colantuoni et al. 2011). Genotypes were called using BeadExpress software. SNPs were removed if the call rate was <98% (mean call rate for this study >99%), if not in HWE (p<0.001) in Caucasian or African American samples, or not polymorphic (MAF<0.01). The total number of observed SNPs remaining in the analysis was 625,439 (96.2%). We then performed genome-wide imputation using the 1000 Genomes reference panel, Shapelt for pre­phasing of haplotypes (Delaneau et al. 2013) and Impute2 software package (Howie et al. 2009). After removing imputing SNPs with MAF < 5% and missingness > 10% within each dataset, there were 6,045,752 SNPs. We identified cis expression quantitative trait loci (eQTLs) using the MatrixEQTL R package (Shabalin 2012), adjusting for race, sex, and surrogate variables after dropping gene expression probes mapping to genes that were not present in the RefSeq database (by gene symbol), and retained eQTLs significant at FDR < 5%.

We then assessed the enrichment of significant DNAm-expression pairs being eQTLs based on identifying the most associated DNAm probe within 50kb of the gene body mapped to each probe, and computing Pearson correlation for each DNAm-expression pair (one statistic per probe). We then used the Wilcoxon rank test comparing test statistics for those probes that were eQTLs compared to probes that were not eQTLs.

### Enrichment for schizophrenia genetic risk

We obtained linkage disequilibrium-clumped results from the latest Psychiatric Genomics Consortium (PGC) genome-wide association study for schizophrenia, which combined datasets used in the PGC1 with data from Sweden (Ripke et al. 2013), which defined genetic regions associated with schizophrenia risk. Enrichment was assessed using the design of the Illumina 450k as background, rather than relying of gene-based models. Using the probe groups from the Illumina 450k, we created 2×2 tables at different p-value thresholds in the SZ GWAS that assessed whether each probe group was in a developmental DMR and/or in a SZ risk region, and quantified this enrichment using a chi-squared test, and report the p-values for enrichment in Table 1. Similar enrichments were calculated for significant loci for IBD, Crohn’s, and UC available in Supplementary Table 2 (LD range provided in first column) of Jostins et al. (2012). Linkage disequilibrium blocks were constructed for all marginally significant T2D loci (p<1x10^−4^) from http://diagram-consortium.org/downloads.html (Stage 1 GWAS Summary Statistics table) using SNAP (Johnson et al. 2008), and collapsed into non-overlapping blocks, retaining the smallest p-value for each LD block.

## Data Access

Data, in both raw and processed forms, will be deposited in the Gene Expression Omnibus (GEO).

